# Heparan Sulfated Glypican-4 is Released from Astrocytes Predominantly by Proteolytic Shedding

**DOI:** 10.1101/2021.02.17.431702

**Authors:** Kevin Huang, Sungjin Park

## Abstract

Astrocytes provide neurons with diffusible factors that promote synapse formation and maturation. In particular, glypican-4/GPC4 released from astrocytes promotes the maturation of excitatory synapses. Unlike other secreted factors, GPC4 contains the C-terminal GPI-anchorage signal. However, the mechanism by which membrane-tethered GPC4 is released from astrocytes is unknown. Using primary astrocyte cultures and a quantitative luciferase-based release assay, we show that GPC4 is expressed on the astrocyte surface exclusively via a GPI-anchorage. Soluble GPC4 is robustly released from the astrocytes predominantly by proteolytic shedding and, to a lesser extent, by GPI-anchor cleavage, but not by vesicular release. Pharmacological, overexpression, and loss of function screens showed that ADAM9 in part mediates the release of GPC4 from astrocytes. The released GPC4 contains the heparan sulfate side chain, suggesting that these release mechanisms provide the active form that promotes synapse maturation and function. Overall, our studies identified the release mechanisms and the major releasing enzyme of GPC4 in astrocytes and will provide insights into understanding how astrocytes regulate synapse formation and maturation.

**Significance Statement:** Astrocyte-derived diffusible factors regulate synapse development and function. However, the regulatory mechanism underlying the release of astrocyte-derived factors is poorly understood. Noting that, unlike many other secreted factors, glypican-4/GPC4 is GPI-anchored, we characterized the release mechanism of GPI-anchored GPC4 from astrocytes and identified the releasing enzyme. Heparan sulfated GPC4 is robustly released from the astrocytes predominantly by proteolytic shedding. In particular, ADAM9 in part mediates the release of GPC4 from astrocytes. Our study provides an enzymatic mechanism for releasing GPC4 from astrocytes and will provide a novel opportunity to understand the regulatory mechanism of neuron-glia communication for synaptogenesis.

## Introduction

Surface shedding is a mechanism by which specific surface proteins are cleaved and an ectodomain is released into the extracellular space. The shedding or release of surface proteins have been found to play important roles in various biological functions (Lichtenthaler et al.). Ectodomain shedding can act to stop activity of proteins at the membrane, such as removing cell surface receptors or cell adhesion proteins, as in the case of N-cadherin (Reiss et al., 2005). Alternatively, shedding can release an active protein from the cell surface to act as a long range, diffusible signaling factor such as syndecan-1 in FGF-2 activation during wound healing (Kato et al., 1998). In this context, shedding can allow for temporally and spatially regulated release or activity of a biomolecule. In other cases, shedding acts as a molecular switch, changing the signaling function of the protein from one role while anchored to the surface membrane to another when shed. For example, membrane-anchored TNF-α can activate TNFR2 upon direct cell to cell contact to activate anti-inflammatory pathways, however when shed, TNF-α acts as a paracrine signal to drive proinflammatory TNFR1 signaling (Grell et al., 1995). In other cases, shed proteins can change the signaling functions of related receptor complexes, such as when released soluble αCNTFR converts the LIF receptor complex and transduces CNTF signaling in the cells that are not normally responsive to CNTF (Davis et al., 1993).

The release of surface proteins also been shown to play important roles in synapse development (Lichtenthaler et al., 2018). For example, Nogo-66 receptor ectodomain shedding drives excitatory synapse formation *in vivo* (Sanz et al., 2018). Proteolytic shedding of cell adhesion molecules causes the structural and functional modifications of synapses (Bajor and Kaczmarek, 2013; Nagappan-Chettiar et al., 2017). Glypican 4/GPC4 has been identified as a cell surface protein sufficient to drive neuronal synaptogenesis *in vitro* (Allen et al., 2012). GPC4 is a GPI-anchored protein (Filmus et al., 2008) and highly expressed in astrocytes (Cahoy et al., 2008; Zhang et al., 2014). Notably, GPC4 expressed in astrocytes are released from the cell and the released GPC4 facilitates synapse maturation in vitro (Allen et al., 2012). While GPC4’s downstream mechanism of action of signaling through RPTP-Δ has been characterized (Farhy-Tselnicker et al., 2017), little is known about how GPC4 is shed or released from astrocytes. A better understanding of the release mechanism of GPC4 offers insights into the synaptogenic role of astrocytes in neuronal development and could lead to novel therapeutic targets for synaptopathic disorders such as autism, schizophrenia, Down syndrome, and epilepsy (Dowling and Allen, 2018; Wilson and Newell-Litwa, 2018).

GPI-anchored proteins can be released from the producing cells via multiple mechanisms, which include proteolytic shedding, GPI-anchor cleavage, and vesicular release (Muller, 2018). There are GPI-anchored proteins which are shed through proteolytic mechanisms such as metalloprotease-mediated shedding of Prion protein and UL16 binding proteins, MHC class I-related molecules (Fernandez-Messina et al., 2010; Linsenmeier et al., 2018). GPI-anchored proteins can also be shed though a lipase-based cleavage mechanism (Park et al., 2013; van Veen et al., 2017). In this context, unlike proteolytic shedding which releases an ectodomain fragment, an entire protein is released from the cell. GPI-anchored proteins have also been shown to associate with a vesicular membrane or lipid particle while becoming separated from the cell (Muller, 2018).

The mechanism by which GPC4 is released has implications to the role GPC4 plays in downstream signaling. GPC4 is post-translationally modified with heparan sulfate side chains at residues near the C-terminus of the protein. These heparan sulfate side chains are necessary for the known synaptogenic signaling functions of GPC4 (Allen et al., 2012; Condomitti et al., 2018). Thus, identification of the release form of GPC4 is critical to understand how astrocyte-derived GPC4 regulates synaptogenesis.

Here, we show that GPC4 is released from the cell surface of astrocytes *in vitro*. We observed that GPC4 is released predominantly via proteolytic shedding, and to a lesser extent by GPI-anchor cleavage, but not by vesicular release. The released GPC4 contains heparan sulfate attachment, suggesting that both release mechanisms preserve the synaptogenic activity of GPC4 in the extracellular space. We also identify ADAM9 as a proteolytic sheddase for GPC4. Overall, these observations show that astrocytes release heparan sulfate containing GPC4 by multiple mechanisms.

## Materials and Methods

### Animals

All animal care and experiments were conducted in accordance with NIH guidelines and the University of Utah IACUC committee (protocol no. 21-02004). C57Bl6/J mouse lines were maintained under the normal housing conditions with food and water available *ad libitum* and 12h light/dark cycle in a dedicated facility at the University of Utah. All primary astrocyte cultures were generated using neonatal wild-type mice of either sex.

### Statistical Analyses

Statistics were performed using Graphpad Prism software version 9. Experiments with two groups were compared with unpaired t-tests and 95 % confidence intervals were calculated for group means and the difference between means. Experiments with three or more groups were compared with ANOVA and Tukey’s multiple comparisons, and 95 % confidence intervals were calculated for group means and the differences between means. One sample t-tests were used for Figure 5A to calculate the difference between control and experimental groups. All data are plotted as mean, with error bars denoting 95 % confidence intervals and all individual data points plotted. For all experiments, the n numbers shown refer to the number of replicates, while N numbers refer to the number of biological replicates.

### Cell Culture

Heterologous cell experiments were conducted using HEK293T cells. Cells were grown in culture media: DMEM (Gibco 11965-092), 10 % FBS (Invitrogen 16140071), 1 % Sodium Pyruvate (Invitrogen 11360070), 1 % Penicillin/Streptomycin (Invitrogen 15140-122). For Biochemistry experiments, cells were seeded between 0.15-0.2 million cells/well in PEI coated 12-well plates. Plasmid constructs (pCAG nHA GPC4, see Table 1) were transfected with Fugene (Promega E2691)/ opti-MEM mixture according to manufacturer protocols. 24 hours after transfection, culture media is changed to serum-free DMEM for biochemical experimental conditions.

**Table 1:**
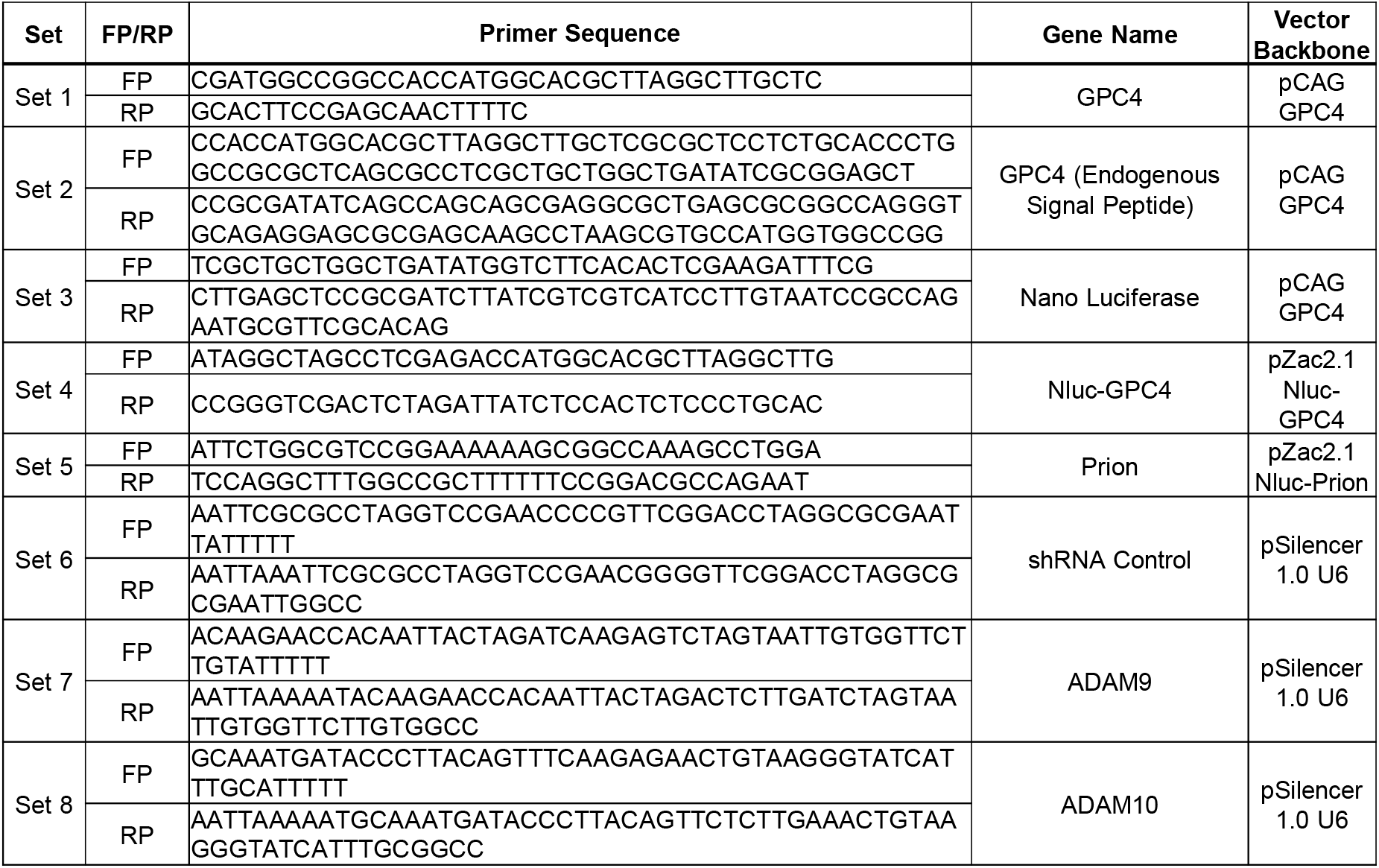
Primers Used

**Table 2:** Statistics

| Data | Data structure (normality test) | Type of test | Power |
| --- | --- | --- | --- |
| Figure 2D: Representative results from one luciferase assay | Yes | N/A |  |
| Figure 2E: Overnight release of GPC4 compared to Prion, normalized to lysate | Yes | Unpaired T-Test | Cohen's D=3.92, P Value <0.0001 |
| Figure 3A: Live surface Biotinylation of GPC4 release over time | Yes | ANOVA; Tukey Multiple Comparisons | F=112.07, P Value <0.0001 |
| Figure 3B: Ultracentrifugation of Nluc-GPC4 in Astrocyte Conditioned Media | Yes | Unpaired T-Test | Cohen's D=7.829, P Value <0.0001 |
| Figure 3C: Pulldown of Nluc-GPC4 by Alpha Toxin | Yes | ANOVA; Tukey Multiple Comparisons | F=206.6, P Value <0.0001 |
| Figure 4A: Reduction of Nluc-GPC4 by Protease Inhibitor GM6001 | Yes | Unpaired T-Test | Cohen's D=1.118, P Value =0.0003 |
| Figure 4B: Reduction of Nluc GPC4 release by ADAM9 shRNA KD | Yes | ANOVA; Tukey Multiple Comparisons | F=35.16, P Value <0.0001 |
| Figure 4C: Induction of Nluc GPC4 release by ADAM9 Overexpression | Yes | ANOVA; Tukey Multiple Comparisons | F=137.1, P Value <0.0001 |
| Figure 5A: Heparinase treatment of Nluc GPC4 prevents pulldown by Anion Exchange Column | Yes | One Sample T-Test | P Value =0.0365, t=5.087, df=2 |
| Figure 5B: PIPLC and Basal released GPC4 show a similar level of binding to Anion Exchange Columns, suggesting both groups release Heparin Sulfate modified GPC4 | Yes | Unpaired T-Test | Cohen's D=.3909, P Value=0.9467 |

Astrocyte culture was generated from P2 mice of either sex. P2 mouse cortexes are dissected and incubated in a Papain solution for 30 mins at 37C. The tissue is then washed with glial media (MEM (Mediatech 15-010-CV), 10 % Horse Serum (Hyclone SH30074.03), 1 % Penicillin/Streptomycin/Glutamine (Invitrogen 10378-016)) before pipette dissociation and filtering through a 70um strainer. Astrocytes are plated on a Poly-D Lysine (Millipore A-003-E)/Collagen (Advanced BioMatrix 5005-100ML) coated T75 flask (Fisher 07-202-000). Astrocytes are grown to confluency, with complete media changes every 3-4 days. Once confluent, astrocytes are harvested with an 8 min TrypLE digestion and resuspended in glial media. Astrocyte cell suspension is nucleofected using the Lonza Nucleofector 4D system at 3 μg plasmid DNA per construct (pZac2.1 Nluc-GPC4, pZac2.1 Nluc-Prion, pSilencer 1.0 U6 shRNA, see Table 1; pCMV3 C-HA ADAM9 (Sino Biological MG50044-CY), pCS2 cHA3 mADAM10 {Park, 2013 #293} with setting CL-133, before being plated on Poly-D Lysine/Collagen coated 12-well plates and allowed to grow to confluency for biochemical experiments. Once confluent, glial media is changed to DMEM high glucose (Invitrogen 11960069) media for biochemical assays and experimental treatments.

### Cloning

GPC4 was cloned from mouse cDNA library from RT PCR and inserted into a pCAG vector (Table 1, Set 1). An HA tag was inserted after GPC4’s signal peptide sequence by long primer (Table 1, Set 2). This construct, pCA N-HA GPC4, was used for HEK293T cell experiments. Nluc-GPC4 was developed by insertion of nanoluciferase (from addgene plasmid #66579) after the cleavage site of the GPC4 signal peptide (Table 1, Set 3), then cloning the resulting construct into a pZac2.1 GFABC1D vector (Addgene 92281) (Table 1, Set 4). Nluc-Prion was cloned from RT PCR of mouse cDNA library and inserted into pZac2.1 GFABC1D vector (Table 1, Set 5). shRNA constructs were designed *in silico*, and cloned into pSilencer 1.0 U6 with long primers (Table 1, Set 6, 7, & 8 for control, *Adam9*, and *Adam10*, respectively).

### Pharmacology

Astrocytes were cultured as described above. Prior to treatment, the media of astrocytes were changed to serum free media and incubated overnight. Astrocytes are then changed into fresh serum free media with and without treatment (25 μM GM6001) and incubated for 3 hours before collection and astrocytes are lysed for biochemical analysis or luciferase assay.

Phosphoinositide phospholipase C (PI-PLC) is a bacterial enzyme that robustly cleaves GPI-anchored proteins from the cell surface membrane. For biochemical assays, PI-PLC is added to cell culture media for 0.5-3hrs for acute treatment or overnight for chronic treatment. Both time frames result in near complete release of all GPI-anchored proteins from the cell surface.

### Western

Media samples were mixed with 4x Gel Loading Buffer (200 mM Tris, 8% SDS, 24% Glycerol, 0.04 % Bromophenol Blue, pH 6.8) while lysate samples were created by lysing cells with 1x Gel Loading Buffer and syringe lysing. Where necessary, astrocyte conditioned media samples were concentrated 20x in 10 kDa cut-off centrifuge protein concentrators (Thermo Fisher 88517) to enrich for WT GPC4 before mixing with gel loading buffer. Samples were loaded into 10 % gels and run at 140 V for 70 minutes, then transferred at 100 V for 60 minutes on PVDF membranes. Membranes were blocked for 1 hour with 5 % milk in TBST (100 mM Tris, 150 mM NaCl, 0.1 % Tween 20, pH 7.5) before placed in primary antibody: HA (BioLegend 901501), GAPDH (Cell Signaling Technology 2118S), Nanoluciferase (R&D Biosystems MAB100261), CD9 (Abcam ab92726). Blots were probed with HRP conjugated secondaries (mouse (Jackson Immuno Research 715-035-151), rabbit (Jackson Immuno Research 705-035-003), rat (Jackson Immuno Research 112-035-003)). Blots were developed using a chemiluminescent substrate (Thermo Scientific 34580).

### Luciferase Assay

Promega Nanoluc Furimazine is diluted 1:50 in Assay Substrate:Assay Buffer, then diluted 1:4 in 1x PBS. Media samples are cleared by centrifugation and 20 μl of sample are loaded in 96 well plate wells. Lysate samples are generated by treatment of 1x Passive Lysis Buffer (Promega E1941) to 12 well plate cultures, and 20 μl of sample are loaded in 96 well plate wells. Luminescence of individual wells was read in a plate reader after injection of 100 μl Furimazine. Average luminescence value is calculated and normalized to either pre-treatment or lysate samples as a transfection control.

### Biotinylation

Cells are cooled on ice, then washed with PBS ++ (PBS, 1 mM CaCl_2_, .5 mM MgCl_2_, pH 7.4) before 10 minute incubation of Sulfo-NHS-SS-biotin (Invitrogen 21331) (1 mg/ml in PBS++). Biotin washed off with PBS++, then quenched with cold glial media, then washed again in PBS before incubated in serum free media for 5 hours at 37C. Incubation media collected, then cleared by centrifugation. 300 μl 1:1 slurry strepavidin beads (Invitrogen 29201) were prepared with a 1x RIPA (150 mM NaCl, 1 % P-40, 0.5 % NaDOC, 0.1 % SDS, 25 mM Tris) wash, two wash buffer rinses (40 mM Tris, pH 7.6) before resuspended in wash buffer as 1:1 slurry. 40 μl 1:1 slurry used per 900 μl media sample and incubated together with rotation at 4C for 1 hour.Beads are spun down, supernatant is collected for analysis, then beads are washed with 1x wash buffer. Bead fraction is resuspended in elution buffer (2 M NaCl, 40 mM Tris, pH 8) and slurry is used for Western and nanoLuciferase assays.

### Ultracentrifugation

Astrocytes are grown after nucleofection in T75 flasks until confluent. Astrocytes are incubated in serum free media for 24 hours before media is collected and subjected to serial centrifugations (10 minutes 300 g, 10 minutes 2000 g, 30 minutes 10,000 g). A 1.5 hour, 100,000 g centrifugation pellets extra cellular vesicles (Konoshenko et al., 2018). The supernatant and pellet are collected for nanoLuciferase assays and western analysis. <u>α–Toxin Pulldown</u>

α-Toxin (AT) plasmid was generously provided by Dr. Yeongjin Hong at the Chonnam National University, Korea (Shin et al., 2006). BL21 cells (Fisher NC9122855) expressing α–toxin are pelleted and resuspended in 0.5x RIPA (75 mM NaCl, 0.5 % P-40, 0.25 % NaDOC, 0.05 % SDS, 12.5 mM Tris) before being sonicated. Cell extract is filtered and incubated with washed TALON beads (Takara 635501) at 4C for 30 minutes. After washing, α-toxin coated TALON beads are resuspended as 1:1 slurry in wash buffer (20 mM Tris, 500 mM NaCl, pH7.9), and 60 μl of slurry is added to 900 μl of media samples and incubated for 1 hour at 25C. Beads are pelleted by centrifugation, supernatant is collected, beads are washed with wash buffer before being resuspended as a 1:1 slurry in wash buffer and used in Nanoluciferase assays.

### DEAE Pulldown

DEAE-Sepharose beads (Sigma DFF100-50ML) are prepared with sequential high and low salt washes (2 M NaCl, 40 mM Tris pH 8.0 and 40 mM Tris pH 7.6, respectively) before resuspended as a 1:1 slurry in wash buffer. Media containing Nluc-GPC4 was added to DEAE 1:1 slurry, and incubated at room temperature with rotation for two hours. Media-DEAE mixture is centrifuged and supernatant is collected while the DEAE pellet is washed once with 10x bed volume wash buffer before wash buffer is aspirated. Nluc-GPC4 is eluted from DEAE beads with high salt (40 mM Tris, 2 M NaCl, pH 8) at room temperature for 30 minutes with mild agitation. Eluted Nluc-GPC4 in elution buffer is collected and used for luciferase assay.

### Heparinase Treatment

Samples containing Nluc-GPC4 were treated with Heparinase II (NEB P0735S) and Heparinase III (NEB P0737S) overnight at 37C, then subject to DEAE pulldown described above, before luciferase activity was assayed. Un-treated samples were similarly incubated at 37C and used as controls for Heparinase treatment for the DEAE pulldown and luciferase assay.

## Results

### GPC4 is expressed on the astrocyte surface via a GPI-anchor

*In silico* analysis indicates that GPC4 contains the N-terminal signal peptide and C-terminal hydrophobic patch and is predicted to be a GPI-anchored protein (Figure 1A). To directly test whether GPC4 is GPI-anchored on the cell surface, we first examined whether GPC4 is released from the cell surface by the application of phosphatidylinositol phospholipase C (PI-PLC), a bacterial GPI-anchor cleaving enzyme (Chen et al., 1998). N-terminal HA-tagged GPC4 was transfected into HEK293T cells and media was collected with and without PI-PLC treatment (Figure 1B). PI-PLC treatment greatly facilitated the release of GPC4, indicating that GPC4 is GPI-anchored on the HEK293T cell surface. Interestingly, GPC4 was detected in the medium by western without PI-PLC treatment, suggesting constitutive release of GPC4 from the HEK293T cells.

**Figure 1.**
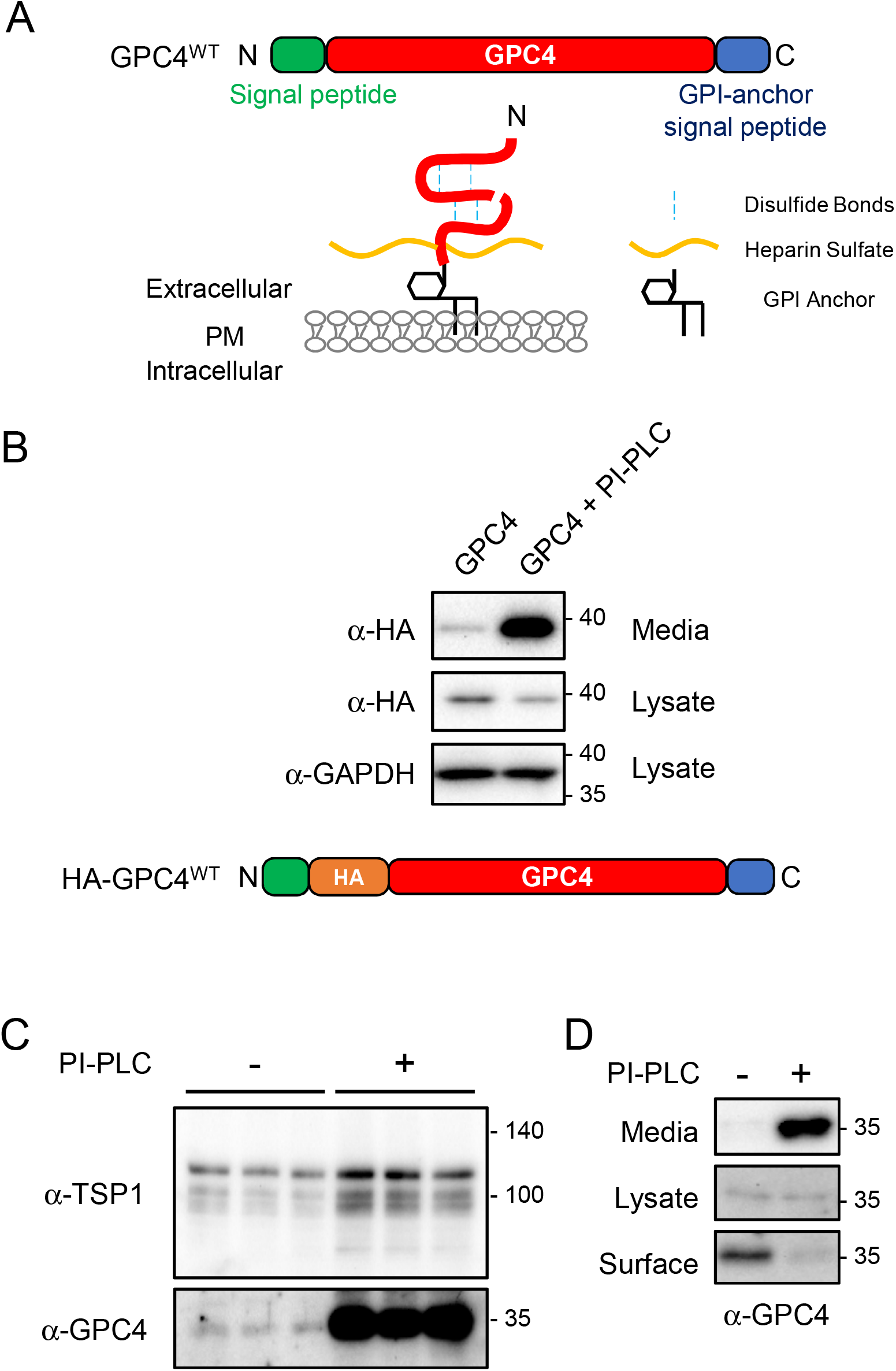
GPC4 is expressed on the astrocyte surface via a GPI-anchorage. ***A***, diagram of GPC4 sequence, protein, and post-translational modifications. GPC4 contains an N-terminal signal peptide and GPI-anchor signal peptide for GPI-anchor attachment. Mature GPC4 is subject to furin cleavage, intrachain di-sulfide bonds, heparan sulfate attachment, and GPI-anchorage to the cell surface membrane. ***B***, Western blot of HA tagged GPC4 construct expressed in HEK293T cells. The ∼37kd band shows the HA tagged N-terminal GPC4 in the reducing condition. PI-PLC treatment in HEK293T cells drives robust release of GPC4, however GPC4 is detected in media in the absence of PI-PLC. ***C***, Western blot of concentrated astrocyte conditioned media for endogenous GPC4 and TSP1, with and without PI-PLC treatment. PI-PLC treatment strongly facilitates release of endogenous GPC4 from astrocytes. ***D***, Biotinylation of surface GPC4 shows that pre-treatment of PI-PLC removes endogenous GPC4 from the cell surface of astrocytes.

Next, we tested whether endogenous GPC4 expressed on the astrocytes is GPI-anchored. Primary astrocytes from mouse cortex were treated with or without PI-PLC overnight and collected astrocyte conditioned media was assayed for shed GPC4. Due to the poor antigenicity of GPC4, we concentrated the astrocyte conditioned medium (ACM) for Western blots (see materials and methods). Consistent with observations in heterologous cells, PI-PLC treatment on astrocytes greatly facilitated the shedding of endogenous GPC4 into the media (Figure 1C). Interestingly, PI-PLC treatment also slightly increased the level of TSP1 in the culture medium, indicating that a portion of secreted TSP1 is associated with GPI-anchored proteins on the astrocyte surface. To determine the degree of GPI-anchorage of GPC4 on the astrocyte cell surface, we performed a surface biotinylation assay, which allows for labeling of all surface proteins at the time of biotinylation. PI-PLC pre-treatment completely removed the biotinylated GPC4 from the cell lysate (Figure 1D). These results indicate that GPC4 is expressed on the astrocyte surface exclusively via GPI-anchor.

### The release of GPC4 from Astrocytes

Proteomic analysis of ACM showed that GPC4 is highly enriched in the astrocyte secretome, ranking 20^th^ in Greco et al. and ranking 28^th^ in our mass spec analysis (not shown) (Greco et al., 2010). However, GPC4 undergoes extensive post translational modifications, including GPI-anchorage, convertase-mediated cleavage, disulfide-linkage, glycosylation, and heparan sulfate attachment (Figure 1A) (Filmus et al., 2008), which may cause the poor antigenicity. To quantify release kinetics of GPC4 from astrocytes, we generated a Nanoluciferease (Nluc) fusion protein construct after the endogenous signal peptide of GPC4 (Nluc-GPC4) (Figure 2A). Nluc is a small (20 kDa) and bright luciferase reacting with a specific substrate, Furamazin (FMZ) (Heise et al., 2013). Attaching the signal peptide to the N-terminus of Nluc efficiently delivers the fusion protein into the secretory pathway. The secreted Nluc shows a linear luminescence in a wide range of Nluc concentration in the medium, which is suitable for quantifying the release of the membrane-tethered protein.

**Figure 2.**
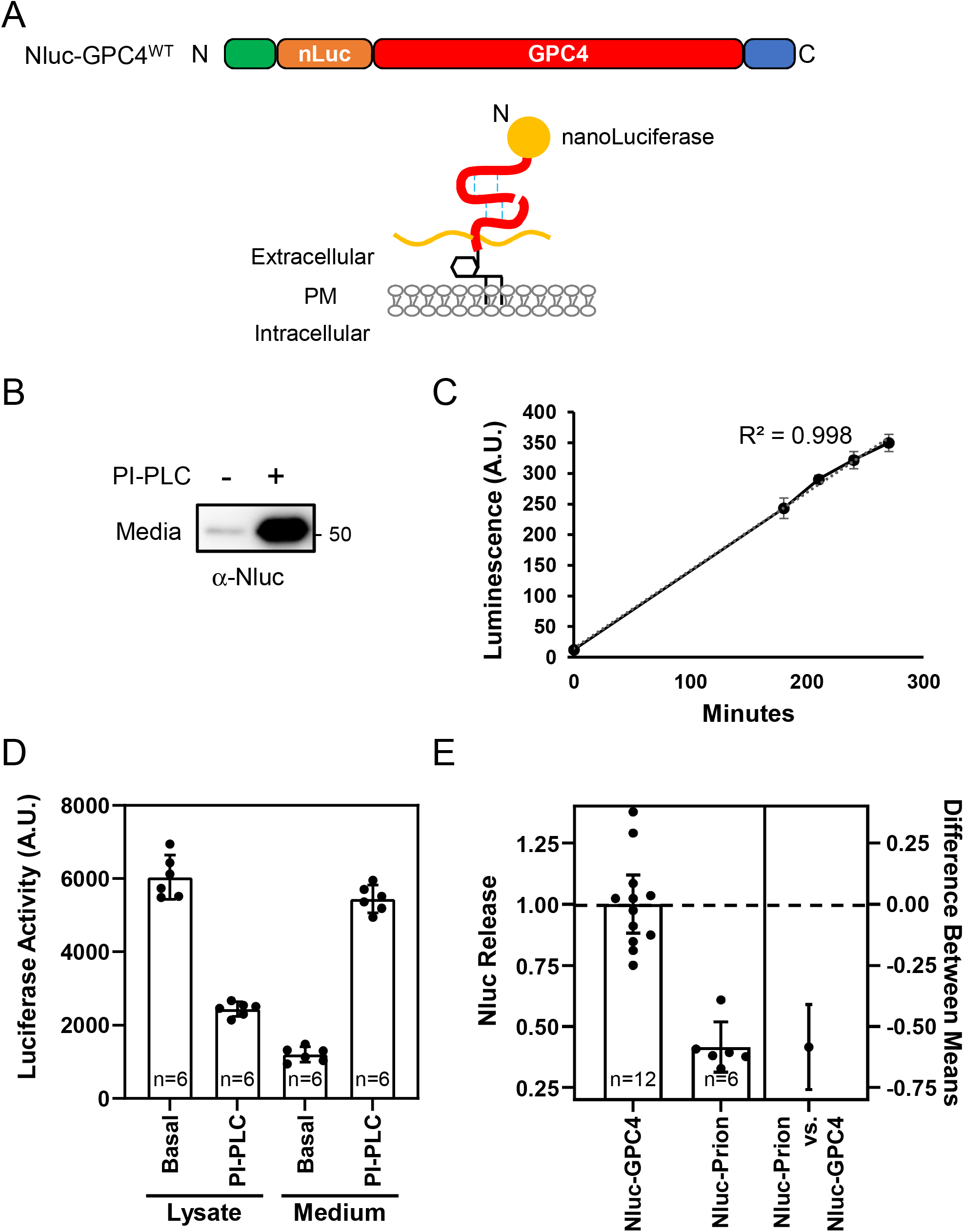
Luciferase assay for quantifying the release of GPC4 from astrocytes. ***A***, NanoLuciferase (Nluc) is inserted at the N-terminus, after the endogenous signal peptide, to preserve GPC4 trafficking. ***B***, Primary astrocytes were nucleofected with Nluc-GPC4 and treated with and without PI-PLC. Western with α-Nluc antibody showed the expected 20kd size shift in the N-terminal fragment. PI-PLC treatment facilitates the release of Nluc-GPC4, confirming the GPI-anchorage of the construct. ***C***, Representative trace of one experiment showing the linear kinetics of GPC4 release from astrocyte culture (R^2^=0.998). Error bars indicate the standard error of the mean. ***D***, Astrocytes expressing Nluc-GPC4 was incubated in a fresh media with and without PI-PLC for three hours and Nluc signal was measured in the cell lysate and media. PI-PLC treatment resulted in the decrease in the luciferase activity of the cell lysate and the corresponding increase in the activity in the media. These data show Nluc-GPC4 is quantitative in measuring released vs. surface pools of GPC4. Error bars indicate 95 % confidence interval of the mean here and in following graphs. ***E***, The release rate (media over lysate activity) of Nluc-GPC4 and Nluc-Prion was normalized to Nluc-GPC4 release. Nluc-GPC4 is released 2-fold faster than Nluc-Prion (T-Test P-value < 0.0001, Cohen’s D=3.92).

We examined the processing, trafficking, and kinetics of the construct as well as the post-translational modifications important for GPC4 function. We nucleofected the Nluc-GPC4 construct under an astrocyte-specific GFABC1D promoter into astrocytes and measured the release of Nluc-GPC4 into the medium. Western blot with Nluc antibody showed the expected ∼20 kDa shift of N-terminal fragment of GPC4 (Figure 2B). Further, the treatment of PI-PLC robustly releases Nluc-GPC4 into the medium, indicating that N-terminal fusion of Nluc does not affect the GPI anchorage and surface expression of GPC4. Interestingly we observed linear accumulation of Nluc-GPC4 signal into the medium at the naïve state (R^2^ = 0.998), further demonstrating the quantitative nature of the Nluc-GPC4 (Figure 2C). Next, we measured the release rate of GPC4 per surface-tethered pool of GPC4. The amount of the surface pool of GPC4 is calculated by the reduction of Nluc-GPC4 signal in the cell lysate and the corresponding increase in the medium signal upon PI-PLC treatment (Figure 2D). We observed that 27 % of the surface GPC4 is released into the medium over 3 hours, which indicates that about 9 % of the surface-expressed pool is released per hour at the naïve state.

Noting the high release rate of GPC4, we compared the release rate of Prion, another GPI-anchored protein abundantly expressed in astrocytes (Cahoy et al., 2008; Zhang et al., 2014). The release rate (luminescence of the medium normalized to that of the cell lysate) of Nluc-GPC4 is about 2-fold higher than that of Nluc-Prion, suggesting that even in this overexpression system, specific release mechanisms exist for different GPI anchored proteins and feature differential release kinetics (Figure 2E) (Mean Difference 95 % CI = -0.7588 to -0.4107, Cohen’s D = 3.92, P-Value < 0.0001). Overall, Nluc-GPC4 retains posttranslational modifications of endogenous GPC4, including furin cleavage, and GPI-anchorage, and allows for rapid, non-destructive measurement of protein release from the cell surface into the media.

### GPC4 is released from the astrocyte cell surface

How is GPI-anchored GPC4 is released from astrocytes? GPI-anchored proteins can be released from the cell via multiple mechanisms, which include (1) secretory pathway bypassing surface expression due to cleavage during processing in the secretory pathway e.g. by GPLD1 (Hettmann et al., 2003; Park et al., 2013), (2) vesicular releases such as in exosomes or lipid associated particles (Dobrowolski et al., 2020; Muller, 2018), (3) GPI-anchor cleavage on the cell surface (Park et al., 2013; van Veen et al., 2017), and (4) proteolytic shedding. Thus we tested each possibility.

We first tested whether extracellular GPC4 at the basal state is shed from the cell surface or directly secreted without being retained on the astrocyte surface membrane. To test this possibility, we conducted a live surface biotinylation assay wherein astrocytes cell surface proteins were biotinylated before being allowed to recover and incubate at 37C for five hours. Incubation media was then collected and assayed for biotinylated GPC4. If GPC4 is secreted directly from secretory pathways, the majority of collected GPC4 should not be biotinylated. Instead we found that the ratio of biotinylated over total released GPC4 is similar between the basal state and the PI-PLC condition, which selectively releases the surface-bound pool (Figure 3A). In contrast, a secreted form of Nluc-GPC4 (GPC4-sec), which lacks the C-terminal GPI-anchorage signal, is secreted without being biotinylated. While we expected the PI-PLC treatment condition to result in a maximal biotinylation ratio, the biotinylation ratio of the basal release was slightly higher than that of PI-PLC conditions (Mean Difference between Basal vs PI-PLC 95 % CI = -0.3189 to -0.1594, Cohen’s D = 1.012, P value = 0.0079). PI-PLC treatment rapidly removes all surface GPC4, which may facilitate the surface incorporation of newly synthesized, un-biotinylated GPC4. This would increase the release of un-biotinylated GPC4 by PI-PLC during the five-hour incubation as compared to untreated control. Together, these data demonstrate that GPC4 is released from the astrocyte surface membrane and not directly from a secretory pathway (Figure 3A).

**Figure 3.**
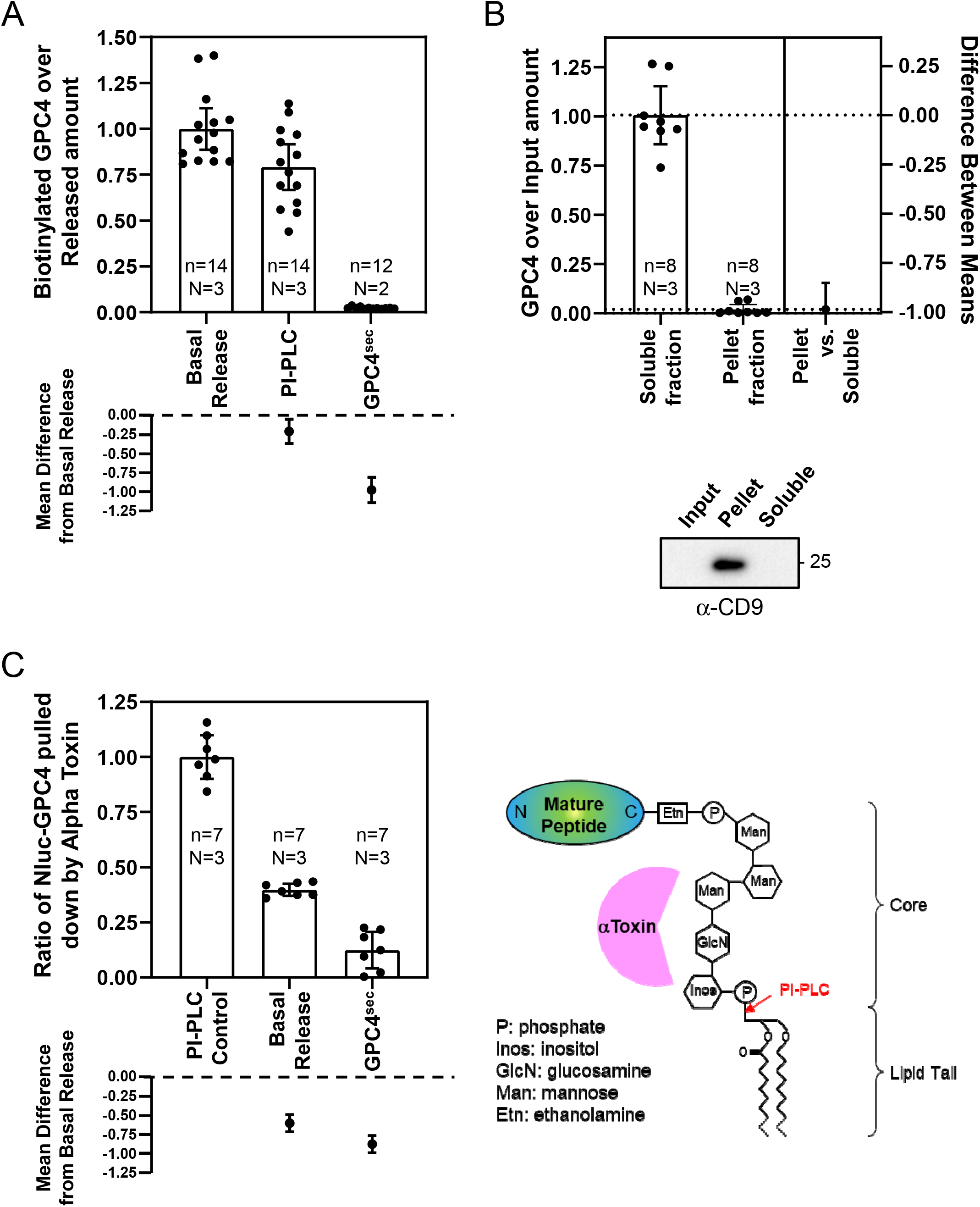
Characterizing the release mechanism of GPC4 using Nluc-GPC4. ***A***, After biotinylating the surface proteins of astrocytes expressing Nluc-GPC4, cells were further incubated in the fresh medium for 5 hours. The released Nluc-GPC4 is collected from the medium and the ratio of biotinylation of the released Nluc-GPC4 was measured (luciferase activity of avidin pull-down over medium input). As a control for the release from the cell surface, PI-PLC were treated during the incubation time. Both basal and PI-PLC induced GPC4 release occur from the cell surface. GPC4^sec^, which does not contain the GPI-anchor signal peptide and is thus not GPI-anchored, is directly secreted without membrane attachment and lacks biotinylation (Basal vs. PI-PLC ANOVA P-value=0.079, Cohen’s D=1.012). ***B***, Ultracentrifugation of astrocyte conditioned media containing Nluc-GPC4 was used to test for GPC4 association with extracellular vesicles as a potential release mechanism. Supernatant (soluble) fraction and pellet (vesicular) fraction Nluc were normalized to input Nluc luminescence. Nluc-GPC4 did not associate with the pellet (vesicular) fraction. An exosome marker, CD9, was enriched in the pellet (vesicular) fraction (T-Test P-value < 0.0001, Cohen’s D=7.829). ***C***, Alpha-toxin, which binds to the glycan core of GPI-anchors, was used to pull down GPI-anchored proteins released through lipase activity. PI-PLC, lipase released Nluc-GPC4 is used as a positive control, while GPC4^sec^, lacking a GPI anchor, is used as a negative control. Nluc-GPC4 pulled down by Alpha-toxin is normalized to input Nluc-GPC4 measurements. Basal release shows partial binding to Alpha-toxin, indicating that ∼30 % of Nluc-GPC4 is released through a lipase mechanism (PI-PLC vs Basal release ANOVA P-value < 0.0001, Cohen’s D 7.659).

### GPC4 is not released through an extracellular vesicle mechanism

Several GPI-anchored proteins are enriched in the extracellular vesicles (Vidal, 2020). These vesicular GPI-anchored proteins retain the lipid tail and are associated with membrane structure. We tested for the possibility that GPC4 is released though an extra vesicular fraction like some GPI anchored proteins such as CNTFR (Choi et al., 2020). In this model of release, GPI-anchored GPC4 is expressed on the cell surface, and released while still bound to the membranes of extracellular vesicles either through exosomal or ectosomal pathways. To test this, media from astrocytes expressing Nluc-GPC4 were sequentially centrifuged, first to separate large particles such as floating cells and cell debris, then to isolate the vesicular fraction by ultracentrifugation (Konoshenko et al., 2018). We measured luciferase activity in the supernatant and pellet fractions, which correspond to soluble protein and vesicle associated fractions, respectively, and normalized the measurements to luciferase activity in the input sample. Luciferase activity shows no change between input and supernatant fractions, with no detection in the vesicular pellet fraction. To validate the ultracentrifugation assay, we tested the pellet fraction and confirmed enrichment of exosomal protein CD9 (Figure 3B). Together, these data show minimal GPC4 presence in vesicular fractions, suggesting that extracellular vesicles are not the major route of GPC4 release in astrocytes.

### GPC4 is partially released though a GPI-anchor lipase

A GPI-anchor lipase such as GDE family enzymes and GPLD1, cleaves the GPI anchor via PLC and PLD mechanism, respectively, and leaves the sugar core attached to the C-terminus of the released protein. In contrast, a proteolytic cleavage would release a GPC4 ectodomain fragment that lacks the GPI-anchor. Noting that α-toxin, a bacterial GPI-anchor binding protein, recognizes the sugar motif of the GPI-anchor (Hong et al., 2002; Shin et al., 2006), we developed a pulldown assay that isolates the species containing the core domain of the GPI-anchor among the released pool (Figure 3C). We expressed α-toxin-His in *E. coli* and purified it with metal resins. To test between these possibilities, we first assayed the percentage of released GPC4 with GPI-anchor residues using a pulldown experiment. Surprisingly, we found that in comparison to our positive PI-PLC control, which releases the surface GPC4 though GPI-anchor lipase activity, constitutively released GPC4 is only partially captured by α-toxin coated beads (Figure 3C) (Mean Difference between PI-PLC control vs Basal Release 95 % CI = -0.4897 to -0.7146, Cohen’s D = 7.659, P value < 0.0001). This suggests that while some GPC4 is released through lipase activity of the GPI anchor, a majority of GPC4 is released through a mechanism that removes the GPI anchor from the released form.

### GPC4 is partially released through a proteolytic mechanism

The results of the α-toxin pulldown immediately suggest proteolytic cleavage as the primary mechanism by which GPC4 is released from the astrocyte cell surface. We first examined the effect of protease inhibitors on GPC4 release. In line with our model of proteolytic cleavage, we observed about 30 % reduction in GPC4 release after treatment with a metalloprotease inhibitor GM6001, an inhibitor of Zn catalyzed proteases, including MMP and ADAM family proteases (Figure 4A) (Mean Difference 95 % CI = -0.5445 to -0.1723, Cohen’s D = 1.118, P value = 0.0003) (Webber et al., 2002).

**Figure 4.**
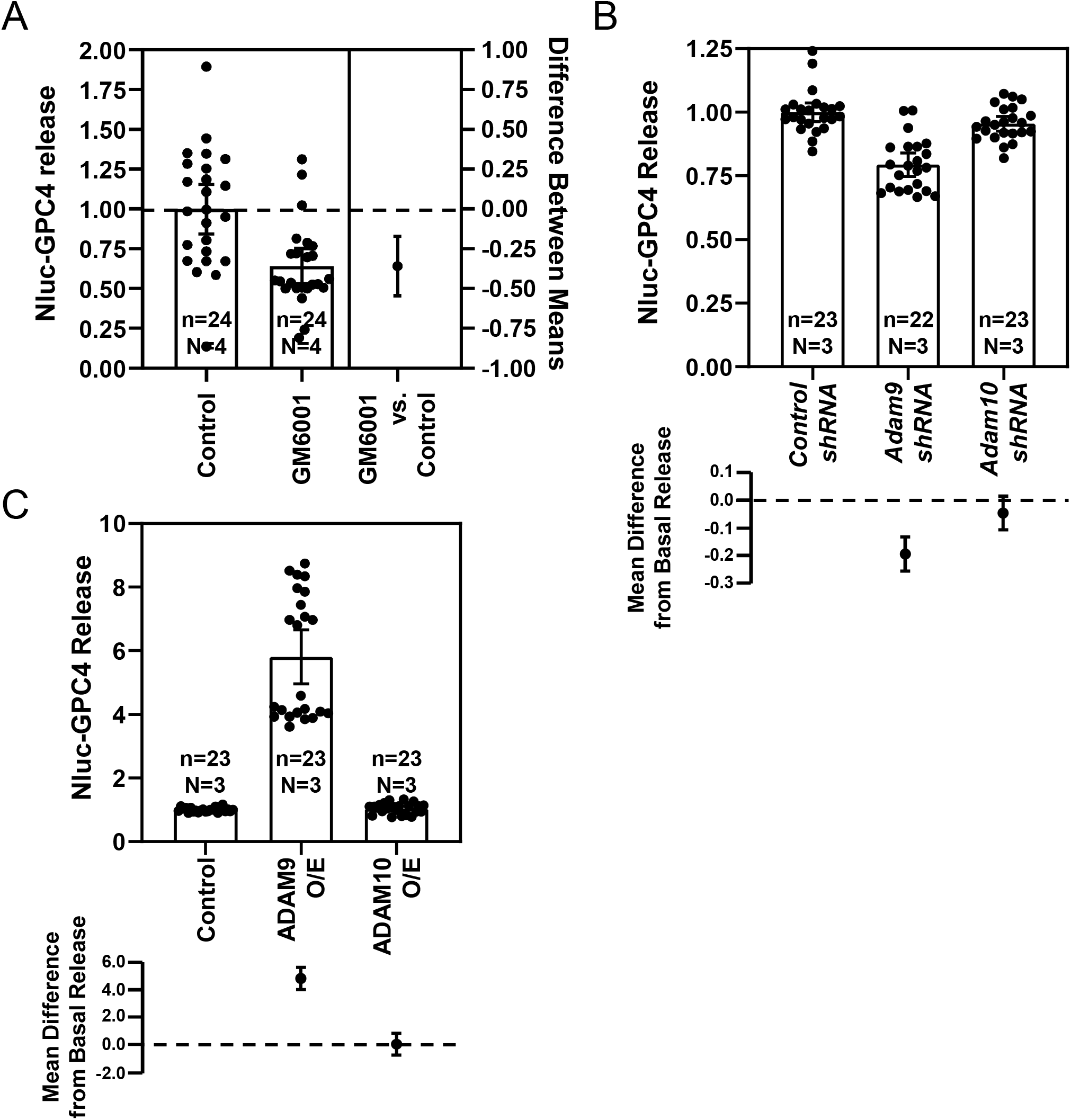
ADAM9 mediates the release of GPC4 from astrocytes. ***A***, GM6001 (25 μM), an inhibitor of Zn^2+^ based proteases including the MMP and ADAM family proteases, reduced the release of GPC4 (T-Test P-value = 0.0003, Cohen’s D = 1.118). ***B***, *shRNA* knockdown of *Adam9* and not *Adam10*, reduces the release of Nluc-GPC4 in astrocytes (Control vs *Adam9 shRNA* ANOVA P-value < 0.0001, Cohen’s D = 2.164). ***C***, ADAM9 overexpression, and not ADAM10 overexpression (O/E), is capable of inducing the release of Nluc-GPC4 from astrocytes (Control vs ADAM9 O/E ANOVA P-value < 0.0001, Cohen’s D = 3.482).

**Figure 5.**
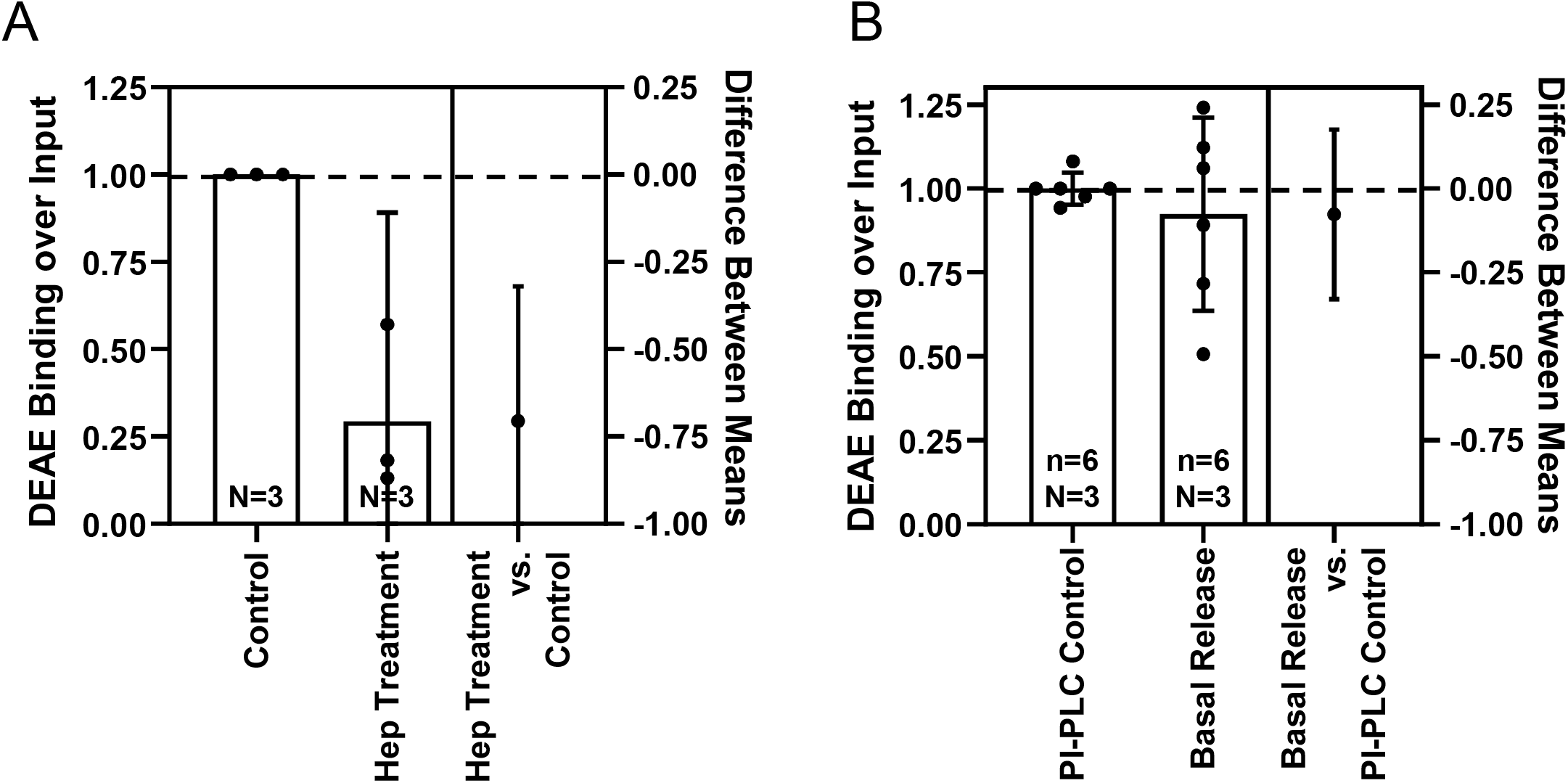
The released Nluc-GPC4 includes the heparan sulfate side chain. ***A***, Heparinase II & III treatment of media Nluc-GPC4 reduced the binding of GPC4 to DEAE anion exchange column, by removing the negatively charged heparan sulfate side chains (One-sample t-test P value = 0.0365, t = 5.087). ***B***, The surface Nluc-GPC4 released by PI-PLC showed similar binding to DEAE as basal release GPC4, indicating that both lipase and protease release mechanisms contain the heparan sulfate modification and are capable of known synaptogenic signaling functions (T-Test P-value = 0.9467, Cohen’s D = 0.390).

### Determining the protease responsible for GPC4 shedding

GM6001 treatment significantly blocked the basal release of GPC4, indicating that Zn-dependent MMPs and ADAM family proteases play a role in GPC4 shedding (Figure 4A). To identify the protease that mediates GPC4 cleavage, we performed a gain of function screen with tissue inhibitors of metalloproteinases (TIMP) which inhibit MMP and ADAM proteases (Brew and Nagase, 2010). Overexpression of TIMP 1 and TIMP 3, which collectively inhibit MMPs and ADAMs 10, 12, 17, 28, and 33, was unable to reduce release of GPC4 from astrocytes (not shown). These results suggest that GPC4 shedding may be mediated by other proteases that are not inhibited by TIMPs, which include ADAM 8, 9, and 19 (Brew and Nagase, 2010). In parallel, we performed shRNA screens of ADAM proteases (*Adamts1, Adamts5, Adam9, Adam10, Adam17*) that are abundantly expressed in astrocytes (not shown) (Li et al., 2019).Among ADAM family proteases we tested, we observed that *shRNA* against *Adam9* significantly reduced the release of GPC4 (Figure 4B) (Mean Difference 95 % CI = -0.1444 to -0.2679, Cohen’s D = 2.164, P value < 0.0001). Consistent with our observations of TIMP overexpression, Adam10 shRNA produced little effect on the release of GPC4 from astrocytes (Figure 4B) (Mean Difference 95 % CI = 0.0154 to -0.1067, Cohen’s D = 0.5948, P value = 0.1807).

To test whether ADAM9 is sufficient to induce the release of GPC4 from astrocytes, we overexpressed ADAM9 together with the Nluc-GPC4 reporter. Our results show that overexpression of ADAM9 and not ADAM10 facilitated GPC4 release from astrocytes (Figure 4C) (ADAM9 Mean Difference 95 % CI = -5.606 to -4.007, Cohen’s D = 3.482, P value < 0.0001; ADAM10 Mean Difference 95 % CI = -0.8448 to 0.7549, Cohen’s D = 0.1542, P value=0.99). Taken together, these data show that ADAM9 mediates release of GPC4 from astrocytes.

### Constitutively shed GPC4 predominantly contains the Heparan Sulfate side chains

Heparan sulfate attachment to GPC4 plays a critical role in downstream signaling, including known synaptogenic functions though RPTP-σ and GPR158 (Condomitti et al., 2018; Farhy-Tselnicker et al., 2017). Unlike GPI-anchor cleavage, which releases full-length protein, proteolytic shedding could generate multiple extracellular fragments. Thus, it is critical to determine whether the released GPC4 contains heparan sulfate attachment sites after proteolytic shedding. To test heparan sulfate attachment to released GPC4, we performed a pulldown assay with an anion exchange column, which binds to the negative charge of heparan sulfate chain (Kojima et al., 1992). As a positive control, we tested whether the surface GPC4 contains heparan sulfate attachment. Treatment of basal condition medium with heparanases reduced the binding of nluc-GPC4 to the column (Figure 5A) (Mean Difference 95 % CI = -1.091 to -0.3206, Cohen’s D = 4.153, One Sample t-test P value = 0.0365), indicating that DEAE binding of GPC4 requires heparan sulfate attachment. Interestingly, the released GPC4 in the basal condition shows a similar binding affinity as that acutely released from the cell surface by PI-PLC (Figure 5B) (Mean Difference 95 % CI = -0.2504 to 0.2181, Cohen’s D =0.3909, P value = 0.9467). These results show that constitutively shed Nluc-GPC4 from astrocytes contains the heparan sulfate modification, indicating that the cleavage site of GPC4 is located downstream of heparan sulfate attachment sites. Overall the released GPC4 from astrocytes by either mode contains the heparan sulfate post translational modification, which is required for its downstream signaling for synapse development.

## Discussion

Here, we show that GPC4 is expressed on the surface membrane exclusively via GPI-anchorage in astrocytes. Our results show that GPC4 is robustly released from the astrocyte surface predominantly by a proteolytic and partially by a lipase shedding mechanism. Soluble GPC4 released by either mode contains the heparan sulfate post translational modification, which is necessary for GPC4 synaptogenic function. Further, our data identifies ADAM9 as a protease that releases GPC4 from astrocytes.

GPC4 was identified as an astrocyte-derived signaling factor. The released GPC4 was shown to induce synapse formation and maturation in cultured neurons (Allen et al., 2012). Further study of astrocyte secreted GPC4 function identified a heparan sulfate-dependent interaction between GPC4 and RPTP-Δ to drive Neuronal Pentraxin 1 (NP1)-mediated recruitment of AMPA receptors (Farhy-Tselnicker et al., 2017). GPC4 is also expressed in neurons and presynaptic GPC4 has been found to play a role in the construction of mossy fiber synapses in the hippocampus (Condomitti et al., 2018). Given the observed roles of GPC4 in both a secreted form in astrocytes and in a cell membrane-tethered form in neurons, the release of GPC4 may play a distinct role in synaptogenesis depending on the cell type. For example, the release of GPC4 from astrocytes may facilitate the maturation of nearby synapses in a non-cell autonomous manner. Alternatively, release of GPC4 from the presynaptic membrane may inhibit the synaptic adhesion of a specific-type of synapse, such as hippocampal mossy fiber synapses. Thus, elucidating the regulatory mechanism of surface GPC4 is critical to understand the physiological roles of GPC4 in synapse development.

To study the mechanism of GPC4 shedding, we found that it was necessary to develop a new tool to counter the poor antigenicity of GPC4. We developed a Nanoluciferease-based GPC4 release assay, which results in a quantitative, sensitive, and non-destructive assay capable of examining the temporal dynamics of GPC4 release. By using this method, we are able to distinguish the different modes of GPC4 release.

Biotinylation assays showed that GPC4 is released from the cell surface, not from the internal pool in the secretory pathway. It is notable that several GPI-anchored proteins are released with a lipid tail-intact form via lipid-associated particles such as on an extracellular vesicle or exosome (Muller, 2018). For example, exosomal GPC1 level is elevated in the serum of patients with pancreatic cancer and is associated with poor prognosis (Melo et al., 2015; Zhou et al., 2018). However, our results show that vesicular release is not a major route of GPC4 release from astrocytes and the majority of released GPC4 exists as a soluble form.

The soluble released form of GPC4 is generated predominantly by proteolytic shedding, which is partially blocked by GM6001, a pan-metalloprotease inhibitor. Our TIMP overexpression and shRNA screens indicated that ADAM9 in part mediates the shedding of GPC4 in astrocytes. However, the magnitude of reduction is limited to 20-30 %, which suggests that there are other proteases working in parallel. Based on our GM6001 results, we expect these proteases to be unaffected by Zinc-dependent proteolytic mechanisms, such as serine proteases like the (TMPRSS) family, which are not inhibited by GM6001 (Lam et al., 2015).

Our data shows that ADAM9 overexpression is capable of shedding GPC4 from astrocytes. It’s possible that ADAM9 induces GPC4 shedding through action on intermediate substrates or activation of an intermediate protease. For example, ADAM9 plays roles in APP shedding through regulation of surface expression and activity of ADAM10 by acting as an ADAM10 sheddase (Moss et al., 2011). We specifically examined the ability of ADAM10 to shed GPC4, and found that neither overexpression nor *shRNA* knockdown of *Adam10* is sufficient to impact GPC4 shedding in astrocytes. Based on these results, we favor the interpretation that GPC4 is a direct substrate of ADAM9, but do not rule out intermediate actors in this shedding mechanism.

α-toxin pulldown assays showed that about 30 % of soluble GPC4 in the medium contains the core structure of the GPI-anchor, indicating a GPI lipase activity on GPC4 in astrocytes. There are several families of GPI-anchor lipases that releases GPI-anchored proteins. The glycerophosphodiester phosphodiesterase (GDPD) family and GPLD1 catalyze phospholipase-C (PLC) and PLD cleavage of the GPI-anchor, respectively (Hettmann et al., 2003; Park et al., 2013; van Veen et al., 2017). Overexpression of GDE2/GDPD5 in neuroblastoma and SH-SY5Y neuronal cells facilitates the release of GPC6 and decreases the surface level of GPC6 (Matas-Rico et al., 2016; Salgado-Polo et al., 2020). Knockdown or knockout of *Gpc6* induces neuronal differentiation similarly to the overexpression of GDE2, suggesting that removal of surface GPC6 by GDE2 mediates the neuronal differentiation in a cell-autonomous manner. Interestingly, a recent study showed that the plasma level of GPLD1 increases after exercise (Horowitz et al., 2020). It is notable that overexpression of GPLD1 wildtype but not a catalytic-dead mutant ameliorates age-related cognitive impairment and enhances adult neurogenesis in aged animals. These results underscore the significant roles of the GPI-anchored proteins and their regulation on the cell surface in neuronal development and function. The GPI lipase that mediates GPC4 release in astrocytes remains to be determined.

The synaptogenic function of GPC4 requires the presence of heparan sulfate side chain. These heparan sulfate side chains are attached to residues proximal to the C-terminus and are thought to mediate binding to GPC4 receptors (Allen et al., 2012; Condomitti et al., 2018; Farhy-Tselnicker et al., 2017). While a lipase shedding mechanism releases GPC4 intact, including the heparan sulfate modification, proteolytic shedding releases a prodomain fragment, and depending on the cleavage site, could shed GPC4 with or without heparan sulfate modification. Our results show that endogenously shed GPC4 is pulled down by a DEAE anion exchange column at similar ratios as PI-PLC lipase shed GPC4, suggesting that endogenously shed GPC4 is heparan sulfate modified and capable of known synaptogenic functions.

Given our results, we expect both lipase and protease shed GPC4 to be capable of functioning as a synaptogenic factor. However, this raises questions as to why there are multiple shedding mechanisms for GPC4 and if the GPI anchor post translational modification plays a role in, or modifies GPC4 function. There could be differences in stability between lipase and protease shed GPC4 in the extracellular space. For example, unlike an ectodomain fragment shed by a protease, the core GPI structure attached to the C-terminus of the protein released by a GPI lipase may play a protective role against extracellular proteases, such as carboxypeptidase E, which is secreted by astrocytes (Ji et al., 2017; Klein et al., 1992). It is also possible that the location and the upstream regulatory signals may differ between proteolytic and GPI lipase enzymes, allowing for differential control of shedding. In addition, given the heterogeneity of astrocyte subpopulations, these activities may be achieved by distinct astrocyte subtypes (John Lin et al., 2017). Future investigations will be required to determine the localization, regulation, subtype-specific expression of the releasing enzymes.

Overall, our study demonstrates that soluble forms of GPC4 are released from the astrocyte surface and contain the heparan sulfate side chain. ADAM9 is a key enzyme that mediates the release of GPC4 from astrocytes. Since, unlike other secreted factors, GPC4 is released from the cell surface by releasing enzymes, our study will provide an opportunity in understanding the regulatory mechanism by which astrocytes promotes synapse maturation and function.

